# Development of neural circuits for social motion perception in schooling fish

**DOI:** 10.1101/2023.10.25.563839

**Authors:** David Zada, Lisanne Schulze, Jo-Hsien Yu, Princess Tarabishi, Julia L Napoli, Matthew Lovett-Barron

**Affiliations:** Department of Neurobiology, School of Biological Sciences. University of California, San Diego. La Jolla, CA, USA 92093

## Abstract

Many animals move in groups, where collective behavior emerges from the interactions amongst individuals. These social interactions produce the coordinated movements of bird flocks and fish schools, but little is known about their developmental emergence and neurobiological foundations. By characterizing the visually-based schooling behavior of the micro glassfish *Danionella cerebrum*, here we found that social development progresses sequentially, with animals first acquiring the ability to aggregate, followed by postural alignment with social partners. This social maturation was accompanied by the development of neural populations in the midbrain and forebrain that were preferentially driven by visual stimuli that resemble the shape and movements of schooling fish. The development of these neural circuits enables the social coordination required for collective movement.

**One-Sentence Summary:** The collective behavior of schooling fish emerges with the development of neural populations selective to social motion.

## Main Text

Animals can navigate their environments as a cohesive group, which provides individuals the benefit of enhanced vigilance and sensing capabilities through interactions with their social partners (*1–4*). Studies of fish schools have demonstrated that collective movement emerges from local interaction rules amongst group members (*5–9*), including short-range avoidance, long-range attraction, and postural alignment (*2*, *4*, *5*). Less is known about how social interaction rules materialize over the course of development (*10*), and the implementation of these rules in the nervous system (*11*, *12*).

Recent studies in zebrafish have demonstrated that fish transition from social avoidance as larvae to social attraction between two and three weeks of age (*13–18*), and that three-week-old zebrafish engage in aggregation behavior through a biological motion-responsive nucleus in the thalamus (*19*). However, zebrafish do not align their postures for extended periods of time (*20*), and it is challenging to image neural activity beyond three weeks of age due to skull ossification throughout development (*21*). Therefore, little is known about the ontogeny of polarization within fish schools, or how developing nervous systems acquire the ability to detect the sensory cues of moving social partners.

Here we address these challenges by studying the collective behavior of the micro glassfish *Danionella cerebrum* (*D. cerebrum*), a recently established vertebrate model system for neuroscience (*22–26*). Their small size, robust social behaviors, and life-long optically accessible nervous system make them a well-suited model organism to address these questions. We report on our investigations of *D. cerebrum* schooling – quantifying the interactions of individual fish, the development of these social interactions, and the maturation of neural populations responsive to schooling-related sensory stimuli.

### Analysis of visually-based schooling in *D. cerebrum* groups

We characterized the collective behavior of *D. cerebrum* adults by recording groups of four fish as they swam around a 300 mm diameter arena (**Fig. 1A, fig. S1A**; see *Methods*) and tracking the position and body orientation of individual fish using SLEAP, a deep-learning based system for multi-animal pose tracking (*27*) (**Fig. 1B, fig. S1B,C**). We quantified the aggregation and alignment of *D. cerebrum* as they explored this open arena, by evaluating the spatial arrangement of fish in groups, the differences in their heading directions, and each fish’s movements in relation to their neighbors (see *Methods*). We assessed these behaviors in both light and dark, to test the role of vision in schooling (*9*, *19*, *22*, *28*, *29*).

**Figure 1.**
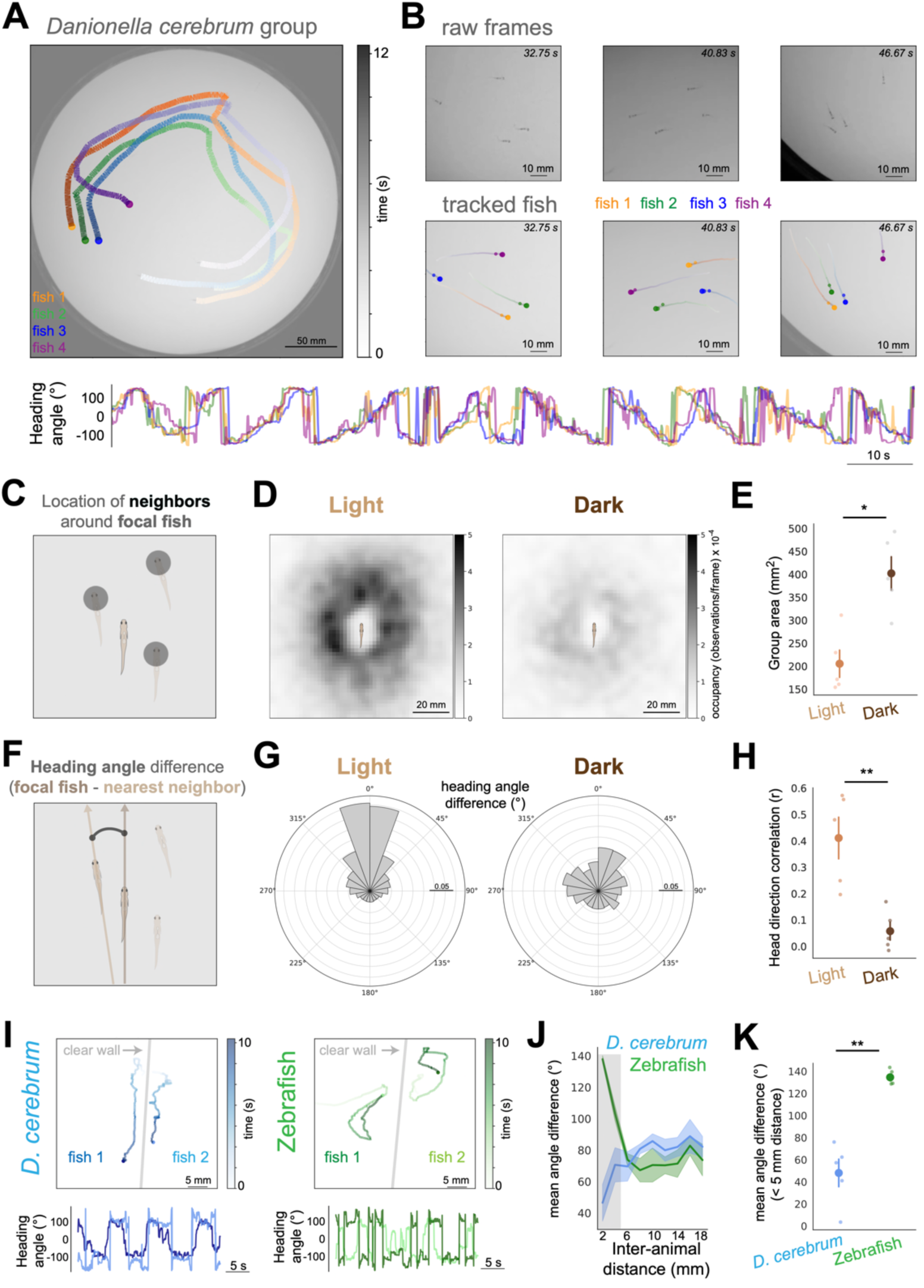
Visually-based schooling in groups of *D. cerebrum.* A) Top: Example of four adult male *D. cerebrum.*, swimming in a 300 mm diameter arena. Projection of tracked movement over 12 s, with each fish shown in a different color. Bottom: timeseries of heading angles of these same fish over two minutes. B) Close-up images of example raw frames (top) and SLEAP-tracking (bottom, with projection of location from prior 1 s) over three example frames. C) Schematic of occupancy measurement: observation of other fish in each frame, relative to the position and heading direction of a focal individual. D) Egocentric occupancy heatmaps, for groups of four fish swimming in light (left), or dark (right). Darker colors indicate more observations of other fish at this position. N = 5 groups, 121 frames/s recordings for 300 s each in light and dark. Average of all timepoints across all fish. Schematic of focal fish is shown (approximate body size). E) Mean group area (mm^2^) comparing between groups of four fish swimming in light versus dark (N = 5 groups). Mann-Whitney U = 1.0. *P* = 0.015. F) Schematic of alignment measurement: difference in the heading angle of the focal individual with its nearest neighbor in each frame. G) Polar histograms of nearest-neighbor angle differences, for groups of four fish swimming in light (left), or dark (right). Observations around 0 °/360 ° indicate aligned neighbors. N = 5 groups, 121 frames/s recordings for 300 s each in light and dark. Histogram of all timepoints across all fish. Scale bar indicates 5% of total observations. H) Mean of pairwise heading direction correlation coefficients (r) amongst all fish, comparing between groups of four fish swimming in light versus dark (N = 5 groups). Mann-Whitney U = 25.0. *P* = 0.00794. I) Example of two adult male *D. cerebrum.* (left, blue) or zebrafish (*Danio rerio*; right, green), swimming on opposite sides of a clear divider. Projection of tracked movement over 10 s. At bottom is a 30 s recording of each animal’s heading direction. J) Difference in heading angle between fish on either side of the clear barrier, with inter-animal distance plotted in 2 mm bins. Shaded area indicates location of close proximity (< 5 mm). Mean +/- s.e.m. K) Mean absolute heading angle difference for fish within 5 mm of each other across the barrier Mann-Whitney U = 0.0. *P* = 0.007936. N=4 pairs each. * *P* < 0.05 \*\**P* < 0.01

We quantified social aggregation by identifying the location of each fish’s social partners relative to itself, and the average size of group distribution in space (**Fig. 1C**). Aggregation was reduced in the dark (**Fig. 1D,E**), with a decrease in the average occupancy surrounding each individual fish and an increase in mean group area (205 ± 29 mm^2^ (mean ± s.e.m) in the light, 401 ± 36 mm^2^ in the dark, **Fig. 1E**; *P*<0.05). In the light, *D. cerebrum* swam faster when in close proximity to their neighbors (**fig. S1D,E**) and executed spatially biased turning behaviors towards conspecifics (**fig. S1F,G**). In addition, each fish moved more slowly (**fig. S1H**; *P*<10^-7^) and less often (**fig. S1I**; *P*<10^-6^) in the dark. We next examined postural alignment by characterizing the difference in heading angle between each fish and its nearest neighbor (**Fig. 1F**). Fish groups were more aligned in the light compared to the dark (**Fig. 1G**), with an decrease in the mean Pearson’s correlation coefficient between group members’ heading in the dark (0.41 ± 0.08 in the light, 0.057 ± 0.03 in the dark; **Fig. 1H**; *P*<0.01). These results demonstrate that vision is necessary for the schooling behavior of *D. cerebrum* (*22*).

Next, we asked if *D. cerebrum* would show aspects of schooling behavior with vision alone (*9*, *28*, *29*). Previous work in juvenile and adult zebrafish has indicated that fish will preferentially engage with conspecifics across a clear barrier (*13*, *30*), where vision is their only source of social sensory input. We conducted similar experiments, comparing adult *D. cerebrum* and zebrafish (**Fig. 1I, fig. S2**, see *Methods*). While closely-spaced pairs of adult zebrafish would orient themselves head-to-head across the clear barrier (*30*) (mean heading angle difference between zebrafish was 134.4 ± 2.9°), *D. cerebrum* pairs would tend towards alignment (mean heading angle difference 47.93 ± 12.45°, **Fig. 1J,K**; *P*<0.01). Thus, *D. cerebrum* exhibit postural alignment with their social partners using vision alone.

### Development of collective schooling behavior

To determine how the key features of schooling behavior – positional aggregation and postural alignment – emerge over the course of development (*31*), we quantified the behavior of *D. cerebrum* groups at two, four, six, and eight weeks of age (**Fig. 2A, fig. SF3A**; see *Methods*), spanning from the larval stage to adulthood. *D. cerebrum* grew in nose-to-tail length from ∼ 4 mm at two weeks to ∼ 12 mm at eight weeks of age, with the densest part of the body identified as the points tracked with SLEAP (**fig. S3B**; see *Methods*). Accordingly, the spatial relationships between fish increased with body length (**fig. S3C-E**), as is expected from interactions based on vision that may use retinal occupancy as an important measure for determining social distance (*7–9*, *17*).

**Figure 2.**
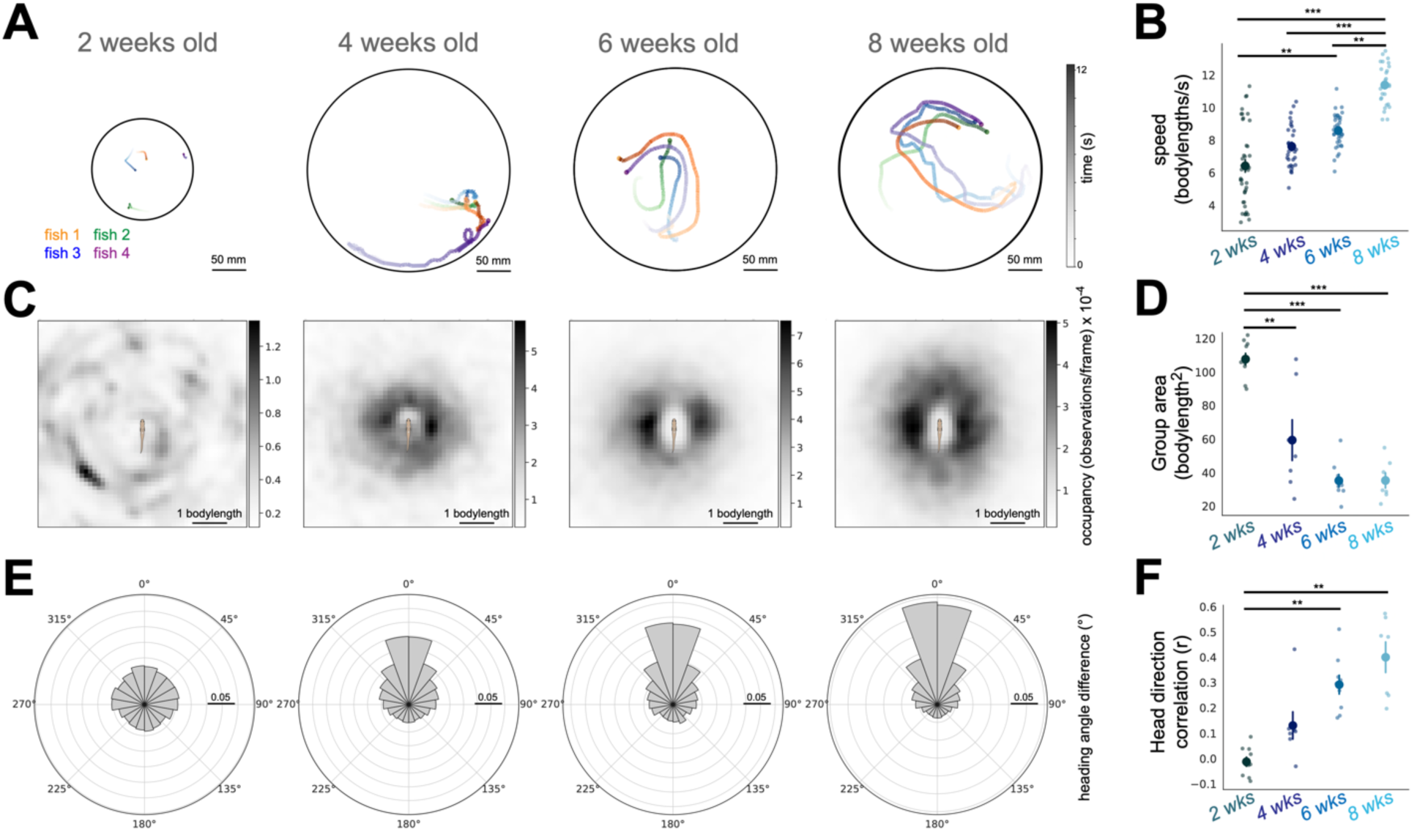
Sequential development of collective behavior in maturing *D. cerebrum.* A) Example of group movement for *D. cerebrum* of different ages (2, 4, 6, or 8 weeks old), swimming in a 300 mm diameter arena (or 150 mm diameter for small 2-week old fish). Projection of tracked movement over 12 s, with each fish shown in a different color. B) Mean speed (body lengths/s) comparing groups of four fish. Kruskal-Wallis followed by *post hoc* Dunn’s test with Bonferroni correction. H = 69.944, *P* = 4.3877 × 10^-15^. C) Egocentric occupancy heatmaps, for groups of four fish (left to right: 2, 4, 6, or 8 weeks old). Maps are scaled by body length, to account for growing size across development. Darker colors indicate more observations of other fish at this position. Average of all timepoints across all fish. Schematic of focal fish is shown (approximate body size). D) Mean group area (mm^2^) divided by mean body length, comparing groups of four fish. One-way ANOVA followed by *post hoc* t-tests with Bonferroni correction. F = 34.477, *P* = 1.072 × 10^-9^. E) Polar histograms of nearest-neighbor angle differences, for groups of four fish (left to right: 2, 4, 6, or 8 weeks old). Observations around 0 °/360 ° indicate aligned neighbors. Histogram of all timepoints across all fish. Scale bar indicates 5% of total observations. F) Mean of pairwise heading direction correlation coefficients (r) amongst all fish, comparing between groups of four fish. Kruskal-Wallis followed by *post hoc* Dunn’s test with Bonferroni correction. H = 23.596, *P* = 3.032 × 10^-5^. N = 10, 7, 9, and 7 groups for 2, 4, 6, and 8 week-old fish, respectively. * *P* < 0.05 \*\**P* < 0.01 \*\*\**P* < 0.0001

We found that fish swam faster as they matured, increasing by 77% from two to eight weeks of age (6.4 ± 0.3 body length/s at two weeks, to 7.6 ± 0.2 body length/s at four weeks, 8.5 ± 0.1 body length/s at six weeks, and 11.3 ± 0.2 body length/s at eight weeks, **Fig. 2B**; *P*<10^-14^). Groups of fish also became more closely spaced with age (**Fig. 2C**), with the mean group area decreasing by 44% between two and four weeks and reaching a 67% decrease by eight weeks (107.4 ± 3.4 body length^2^ at two weeks, to 59.1 ± 12.1 body length^2^ at four weeks, 35.0 ± 3.3 body length^2^ at six weeks, and 35.1 ± 4.5 body length^2^ at eight weeks, **Fig. 2D**; *P*<10^-8^). Accordingly, pairwise distances between individuals in these groups also decreased over development (**fig. S3F**). Therefore, social aggregation in *D. cerebrum* largely develops between two and four weeks of age, and plateaus by six weeks.

In contrast to the rapid development of aggregation, postural alignment between partners developed more slowly (**Fig. 2E**), with progressive increases in the heading angle correlation between four, six, and eight weeks of age (mean Pearson’s r = -0.01 ± 0.02 at two weeks, 0.129 ± 0.05 at four weeks, 0.291 ± 0.03 at six weeks, and 0.399 ± 0.06 at eight weeks, **Fig. 2F**; *P*<10^-4^). This developmental sequence – with the onset of aggregation preceding alignment – was also apparent when sorting fish by speed instead of chronological age (**fig. S3G,H**), and when comparing aggregation and alignment within groups (**fig. S3I**).

To measure this sequential emergence of schooling behavior at higher temporal resolution, we conducted a longitudinal measurement by sampling the same group of fish every 3-4 days from two to six weeks of age. We observed a similar pattern, that group area began to decrease between the ages of 25 and 32 days old, while heading direction correlation increased between 32 and 39 day old fish (**fig. S4**). These results indicate that *D. cerebrum* schooling develops in a sequence – beginning with social avoidance at two weeks of age, then gaining aggregation at four weeks of age and alignment at six to eight weeks of age.

### Imaging brain-wide neural activity in *D. cerebrum* viewing social motion stimuli

As animals develop from social avoidance to aggregation and alignment, their changing behavioral responses to social partners suggests a change in the sensory encoding of conspecific actions (*11*, *12*). We therefore explored how the brains of *D. cerebrum* respond to naturalistic schooling-like visual stimuli, and how these responses mature over social development. To record neural activity from behaving *D. cerebrum*, we developed a novel method for head-tethering and visual stimulus display for behaving animals undergoing large-scale two-photon calcium imaging (**fig. S5A**; see *Methods*). This preparation allows for multi-region cellular-resolution calcium imaging from transgenic *D. cerebrum* with near-pan-neuronal expression of a genetically-encoded calcium indicator (*Tg(elavl3:H2B-GCaMP6s); 25*, *26*). This preparation allows the mouth and gills to be free for respiration, and the tail to be free for behavioral monitoring (see *Methods*). We confirmed that tethered *D. cerebrum* can see visual motion stimuli under these conditions, and identified consistent neuronal responses to widefield optic flow stimuli across ages (**fig. S5B-F**).

In order to create visual stimuli with naturalistic biological motion (*15*, *19*, *32*), we transformed a recording of four schooling *D. cerebrum* into one animal’s egocentric view of their three group members (**Fig. 3A**; see *Methods*). We extracted a short epoch from this egocentric view to display to tethered *D. cerebrum* (**Fig. 3B**), with trials showing these “virtual group members” as either a natural fish-like shape (elongated in the horizontal axis), a sphere, or an inverted fish-like shape (elongated in the vertical axis). We also generated a background social motion stimulus by applying the motion of one virtual group member to the floor underneath the fish (**Fig. 3B**; see *Methods*).

**Figure 3.**
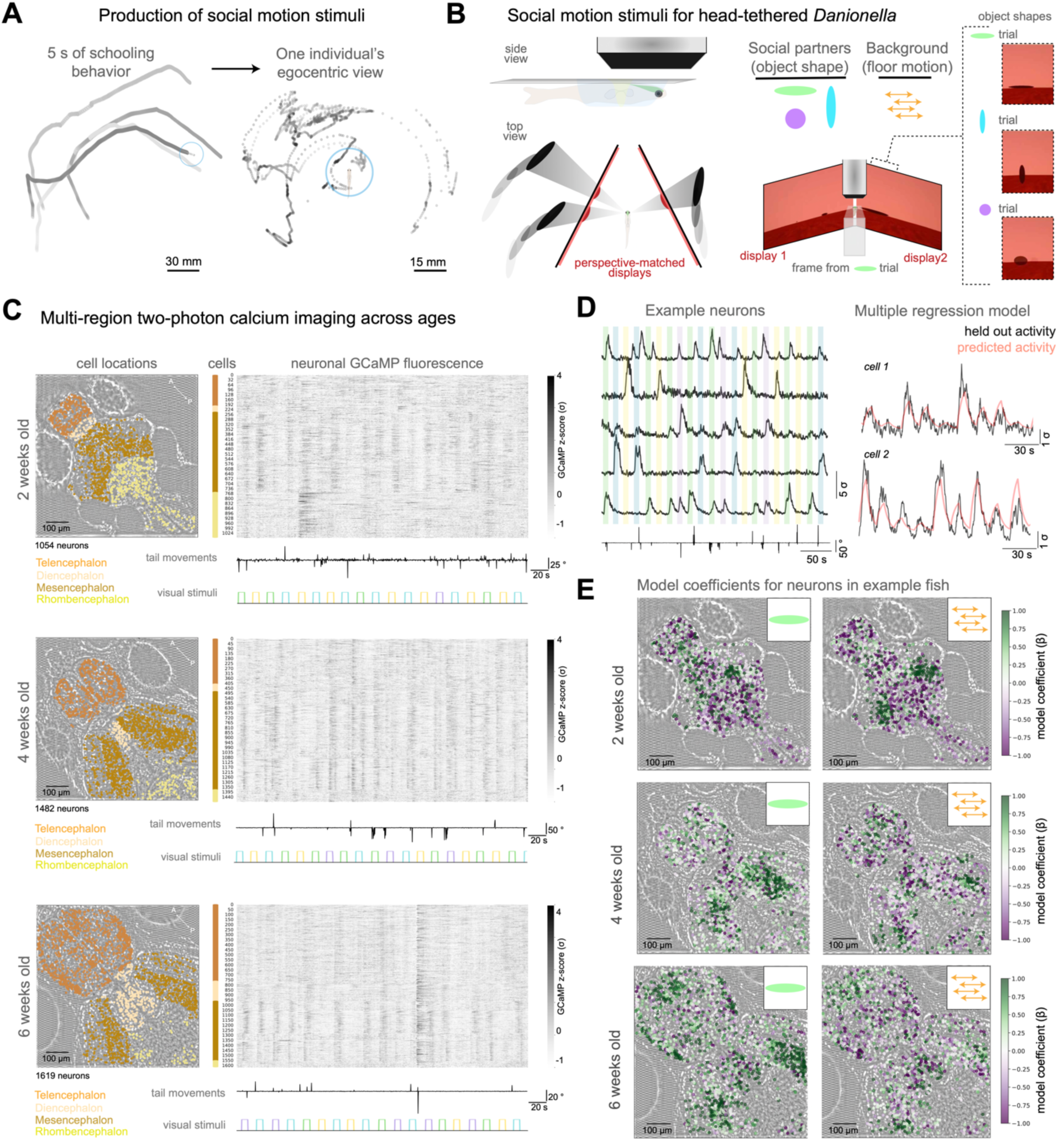
Imaging neural dynamics evoked by social motion across development. A) Production of social motion stimuli for head-tethered fish. A 5 s period of schooling in a group of four 8 week-old fish (left) is rotated around one focal fish to produce this fish’s egocentric view of the other three animals over the 5 s period (right). B) 2, 4, or 6-week old *D. cerebrum* are head-tethered under a two-photon microscope objective with mouth and tail free, and immersed in a visual display environment that covers ∼260 ° of visual space (left). The egocentric social motion stimulus is presented to the fish, where the three virtual group member motions are applied to three varieties of dark shapes (horizontal, vertical, or circular), or to the floor below the fish (“background”). C) Example fish of each age, with anatomically-annotated cell locations (left; A=Anterior; P=Posterior), and z-scored neural activity for all recorded neurons, sorted by anatomical location. Social motion stimulus timing (colors corresponding to the stimulus types in panel B) and tail movements are noted below each neural activity heatmap. A subset of the full imaging time is displayed. D) Left: short epoch of a fish’s example neurons’ responses to social motion stimulus types and tail movements. Color corresponds to the stimuli in panel B. Right: two example cells, with measured z-scored fluorescence (black, final 25% of activity trace) and the predicted fluorescence (red) from a linear model trained on the first 75% of the activity trace (ridge regression trained with 5 regressors – 4 visual motion types and tail movements convolved with a GCaMP-like decay). E) Location of neurons in example fish of each age, with neurons colored by model coefficients (β) for the horizontal shape motion (left) or background motion (right).

We imaged neural activity at cellular resolution across multiple brain regions in *D. cerebrum* immersed in this visual environment, where social motion stimuli were provided for 7 s trials, interleaved with a 9-11 s inter-trial interval; stimuli for each trial were randomly selected to be virtual group members with either horizontal, vertical, or spherical shapes, or background motion (**Fig. 3C**). Tethered *D. cerebrum* executed tail movements throughout, with two-week old fish swimming at high baseline rates (*25*), but with social motion-driven tail movements increasing with age (**fig S6A**). We recorded 3424 cells in two-week old animals (N=4 fish), 3268 cells in four-week old animals (N=4 fish), and 4072 cells in six-week old animals (N=4 fish). Amongst these, many neurons showed increases in fluorescence (>2.5 σ) during the social motion of one or more group member shapes (8% of cells in two week old fish, 14% in four week old fish, 11% in six week old fish, see *Methods*). We sorted neurons into broad anatomical regions (**Fig. 3C, fig. S6B**), and examined the relationship between social motion stimuli and neural activity by fitting a linear model to each neuron with the four visual stimulus types and tail movement as regressors (**Fig. 3D**; see *Methods*). We found that average model coefficients (β) for social motion stimuli increased across development for neurons in multiple brain regions (**Fig. 3E, fig. S6C**). Therefore, neurons are responsive to social motion stimuli even in young larvae, but the fraction of neurons responding and their magnitudes increase over development.

### Maturation of neural populations selective to the shape of social partners

We next asked whether these neurons were responsive to all social motion stimuli, or could distinguish between different classes of stimuli. We analyzed the trial-averaged trajectories of neural activity in low-dimensional space (**Fig. 4A, fig. S7A**; see *Methods*), to determine if these trajectories differ amongst social motion stimuli with virtual group members of different shapes. We first compared the average pairwise distance between trajectories for social motion stimuli with horizontal, vertical, or spherical group members; the average cosine (angular) distance between trajectories did not change in two-week old fish (-0.01 ± 0.04), but increased in four- (0.33 ± 0.08) and six- (0.30 ± 0.12) week old fish (**Fig. 4B**; *P*<0.05). In contrast, the average pairwise distance between trajectories for each shape stimulus and the background motion increased across all groups, with no differences between ages (two-week old fish = 0.77 ± 0.21, four-weeks = 0.72± 0.17, six-weeks = 0.74 ± 0.24; **Fig. 4C**; *P*=0.98). These findings persisted when neurons were subsampled to common numbers of cells across ages (**fig. S7B**, see *Methods*). For both stimulus type comparisons, the Euclidean distance increased between trial-averaged trajectories across all groups, and was not different between ages (**fig. S7C**). Therefore, in socially-mature fish, neural population activity evoked by social motion stimuli follows distinct trajectories depending on the shape of observed group members.

**Figure 4.**
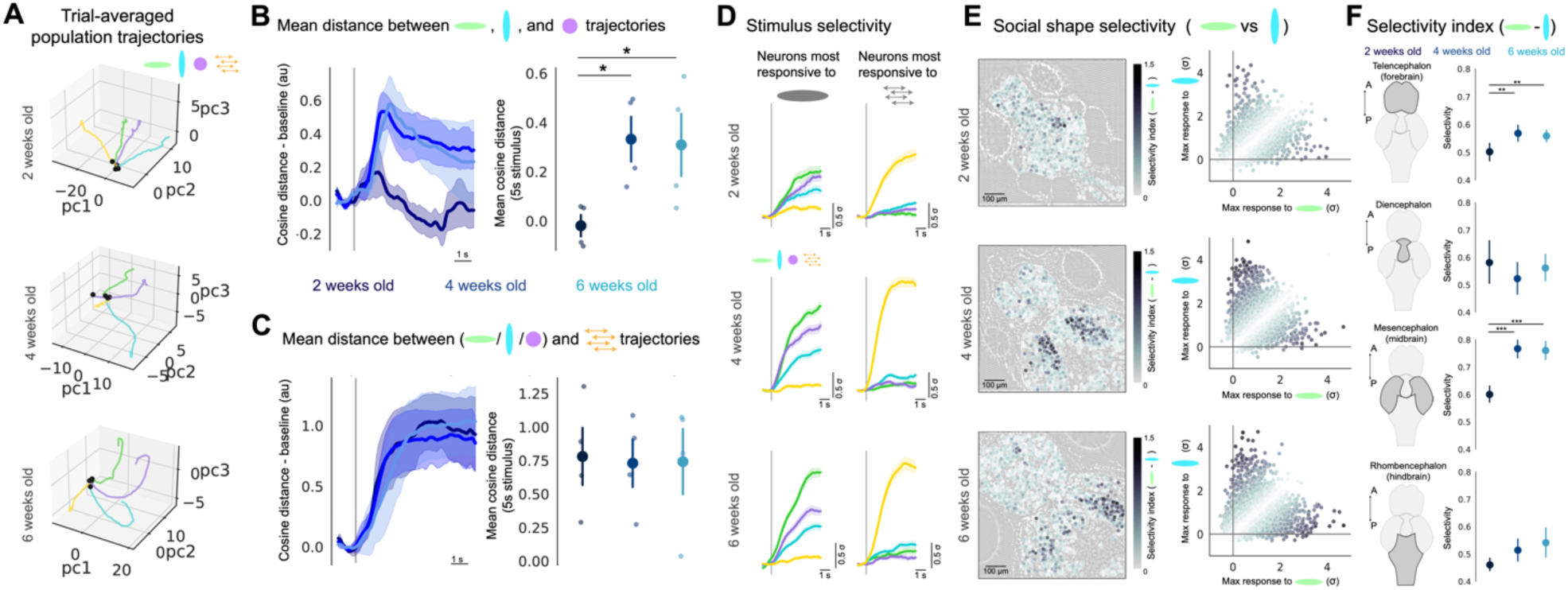
Maturation of neural selectivity to the shape of social motion stimuli. A) Low-dimensional trajectory of trial-averaged social motion stimulus responses, beginning at stimulus onset (black dot) and progressing for 7 s after stimulus onset. Trajectories are colored by stimulus type, and are shown for the first three principal components (pcs). Example fish of each age are shown. B) Mean pairwise cosine distance of trial-averaged trajectories in 15-dimensional PC space, between each of the three virtual group member social motion trial types (horizontal, vertical, or spherical shape). Left: Timeseries are baseline-subtracted (1 s before stimulus onset), and mean +/- s.e.m is shown. Right: mean across the 7 s stimulus period. One-way ANOVA followed by *post hoc* t-test tests with Bonferroni correction. F = 4.436, *P* = 0.045. C) Mean pairwise cosine distance of trial-averaged trajectories in 15-dimensional PC space, between each of the three virtual group member social motion trial types and the background social motion trials. Left: Timeseries are baseline-subtracted (1 s before stimulus onset), and mean +/- s.e.m is shown. Right: mean across the 7 s stimulus period. One-way ANOVA F = 0.015, *P* = 0.984. D) Midbrain neurons for fish of each age were selected as top 5^th^ percentile of model coefficients (β) for horizontal shape social motion (left) or the background social motion (right), and their trial-averaged responses to each stimulus type is displayed. Traces are mean +/- s.e.m, colored by stimulus type, and the stimulus onset is noted by the gray line. E) Object-shape selectivity (absolute value of the difference in the maximum response to horizontal and vertical shape stimuli) across each age, displayed on cell locations in example fish (left) and on a scatter plot of stimulus responses across all fish (right; N= 3424 (Wk2), 3268 (Wk4), and 4072 (Wk6) cells.). F) Summary of shape selectivity measure (absolute value of the difference in the maximum response to horizontal and vertical shapes) across each age, and separated by anatomical subdivision (left; A=Anterior; P=Posterior). Mean +/- 95% confidence intervals are shown. Kruskal-Wallis followed by *post hoc* Dunn’s test with Bonferroni correction. From top: Telencephalon: N = 734 (Wk2), 1016 (Wk4), 1669 (Wk6) cells, H = 11.257, *P* = 0.00359. Diencephalon: N = 130 (Wk2), 199 (Wk4), 435 (Wk6) cells, H = 1.269, *P* = 0.530. Mesencephalon: N = 1580 (Wk2), 1672 (Wk4), 1726 (Wk6) cells, H = 54.537, *P* = 1.437 × 10^-12^. Rhombencephalon: N = 980 (Wk2), 381 (Wk4), 242 (Wk6) cells, H = 6.597, *P* = 0.037. N=4 fish for all groups. * *P* < 0.05 \*\**P* < 0.01 \*\*\**P* < 10^-6^.

We next asked whether these population-level distinctions between neural responses to the shape of moving social partners were a consequence of shape-tuning in single neurons. To determine whether neurons are selective for each class of social motion stimulus, we plotted the trial-averaged activity of the most responsive neurons in each annotated brain region (top fifth percentile of β distribution; **fig. S8**; see *Methods*). We noted particularly robust responses in the midbrain optic tectum, consistent with this structure’s role in detecting ethologically-relevant object motion across species (*33–35*). We analyzed the most responsive midbrain neurons for the horizontal fish-like shape and the background motion stimulus; while responses to different social motion shapes were similar in young fish, they appeared more distinct in older fish (**Fig. 4D**). In contrast, neurons responsive to background motion were selective across all ages (**Fig. 4D**), consistent with the mature retinotopy of motion stimuli even in larval fish (*36*).

We quantified social motion shape selectivity by comparing the maximum responses of each neuron to social motion of the horizontal and vertical shapes (**Fig. 4E**; see *Methods*). We sorted neurons by brain region, and found an age-dependent increase in the mean shape selectivity of neurons in the forebrain (mean response difference of 0.50 ± 0.02 σ at two weeks, 0.56± 0.01 σ at four weeks, 0.55 ± 0.01 σ at six weeks; **Fig. 4F**; *P*<0.005), and midbrain (mean response difference of 0.59 ± 0.02 σ at two weeks, 0.77± 0.02 σ at four weeks, 0.76 ± 0.02 σ at six weeks ; *P*<10^-11^). A similar selectivity metric applied to the comparison of horizontal shapes and background motion also increased with age (**fig. S9**). These data demonstrate that the selectivity of forebrain and midbrain neurons to the shape of schooling group members matures with the social development of *D. cerebrum*.

## Discussion

Collective behavior emerges from the interactions between individuals in a group, that continually sense each other’s actions and move cooperatively. These interactions are produced by the sensory, integrative, and motor processes of each animal’s nervous system – a complex process that arises over the course of development. Here we have used *D. cerebrum* as a model system (*22–25*) to investigate the development of visually-based schooling behavior and its neurobiological underpinnings. We found that *D. cerebrum* schooling behavior is based on vision (confirming prior work (*22*)), and found that these fish can engage in social behavior based on vision alone, similar to zebrafish (*13*, *15*, *30*). We then demonstrated that schooling develops in a sequential manner, where two week-old larvae begin with social avoidance, and progressively acquire the ability to aggregate beginning at four weeks, followed by postural alignment and coordinated swimming beginning at six weeks of age. Future studies can determine whether this developmental progression is innate, or requires social or specific visual experience at some or all stages of this maturation (*12*, *28*).

We established a novel head-tethering method to image neural activity from behaving *D. cerebrum* across developmental stages, and recorded neural activity across brain regions with cellular resolution while presenting visual stimuli that mimic the egocentric experience of a schooling fish. In agreement with previous work in juvenile zebrafish (*19*), we found biological motion-driven neural activity at all ages. However, we found that selectivity to the shape of social motion stimuli develops with age, and is only apparent in older fish. These results indicate that, as *D. cerebrum* matures to engage both the aggregation and alignment aspects of schooling, their nervous systems acquire the ability to identify the shape and posture of moving conspecifics.

While prior studies have suggested that postural alignment in fish schools is a consequence of close aggregation (*6*, *18*), our results suggest that alignment is a separate process. We observe that four-week old fish aggregate without strong alignment, whereas older fish do both. We therefore hypothesize that animals actively engage in alignment, and that this process is enabled by the development of neural circuits that identify the postures of moving social partners. Additional studies are required to determine whether social shape-selective neurons distinguish between other features of conspecifics and their motion (including color, patterning, or sex (*28*)), and their relationship to neurons selective to biological motion (*19*). These mechanisms may be distinct; in zebrafish, knockout of the oxytocin receptor disrupts behavioral preference for biological motion, but not preference for fish-like shapes (*32*).

A central question to address in future studies is how visual representations of conspecifics and their actions are transformed into the movement commands that produce social aggregation and alignment in schooling fish. We hypothesize that these behaviors involve coordination of brain-wide circuits, including the visually-driven midbrain and forebrain neurons we describe, as well as brainstem motor control pathways (*37*) and ascending hypothalamic and neuromodulatory influences (*38*). These studies will benefit from simultaneously imaging all neurons in adult *D. cerebrum*, using newly-developed microscopy techniques (*26*) that can allow for investigation of dynamics across the entire vertebrate social decision-making network (*39*). Investigating the neurobiological mechanisms of collective behavior can provide links across biological scales – from the microscale properties of individual nervous systems to the emergent properties of animal collectives.

## Acknowledgments

We thank Benjamin Judkewitz and Adam Douglass for sharing *Danionella* and advice on their care, as well as Talmo Pereira and the Pereira lab for assistance with animal tracking. We thank Takaki Komiyama, Johnatan Aljadeff, Byungkook Lim, Ashley Juavinett, Scott Sternson, and Priya Rajasethupathy for comments on the manuscript, and thank all members of the Lovett-Barron Lab for discussion, support, and feedback.

## Funding

Human Frontier Science Program Postdoctoral Fellowship (DZ)

Zuckerman STEM Program Israeli Postdoctoral Fellowship (DZ)

Kavli Institute for Brain and Mind Postdoctoral Fellowship 2022140 (LS)

UC San Diego J. Yang Scholarship (JY)

Taiwanese Government Scholarship to Study Abroad Award (JY)

NIH T32GM133351 (JLN)

## Searle Scholars Award (MLB)

Packard Foundation Fellowship (MLB)

Pew Biomedical Scholar Award (MLB)

Klingenstein-Simons Fellowship in Neuroscience (MLB)

Sloan Research Fellowship (MLB)

NIH R00MH112840 (MLB)

NIH New Innovator Award DP2EY036251 (MLB)

## Author contributions

DZ, LS, JY, and MLB designed experiments. DZ, JY, JN, and PT performed freely-swimming behavioral experiments. DZ, LS, JY, PT, JN, and MLB wrote code and analyzed data. DZ, LS, JY, JN, and MLB developed the head-tethering approach. DZ, LS, and MLB performed imaging experiments. DZ, LS, and MLB wrote the paper, with feedback from all authors.

## Data and materials availability

Data, code, and materials will be made available upon publication.

## Supplementary Materials

### Materials and Methods

#### Fish husbandry

*D. cerebrum* and zebrafish were raised and maintained, separately, in standard zebrafish housing systems (Aquaneering; system water temperature 29 ± 0.5 °C, pH 7.0, conductivity 650 mS), under 14 h light/10 h dark cycles, and fed twice a day. *D. cerebrum* were bred in communal tanks of 20–40 individuals with ∼5 cm silicone tubes as spawning environments (*22*, *40*). Eggs were collected during the first hour of daylight, and during 1-2 h after morning and afternoon feeding. Embryos (0-5 days old) were raised in egg-water (0.2 mg/L Instant Ocean, 3 g/l mm CHNaO3, and 0.15% methylene blue dissolved in reverse osmosis purified water) in an incubator at 29 ± 0.5 °C. Larvae (5-14 days old) were co-cultured with L-type rotifers (*Brachionus plicatilis*) in static tanks, where water levels were raised approximately 1 L per day. After entering water circulation at 14 days old, *D. cerebrum* were fed increasingly with artemia. *D. cerebrum* reaches adulthood and lay eggs by 8-10 weeks. Wild-type AB zebrafish were raised using standard procedures, and used for behavior experiments at 6-8 weeks of age. Wild-type *D. cerebrum* were provided by Dr. Adam Douglass (University of Utah) and Dr. Benjamin Judkewitz (Charité Berlin). The transgenic line *Tg(elavl3:H2B-GCaMP6s)* was provided by Dr. Benjamin Judkewitz (Charité Berlin).

#### Behavioral experiments

All behavioral experiments were controlled with BonsaiRx (bonsai-rx.org), using BonVision and BonZeb packages (*41–43*), and custom routines to control and time-stamp camera acquisition and visual stimuli. All behavioral experiments were conducted in a behavioral room at 28 °C maintained with central circulation and portable electric heater. Since the onset of *D. cerebrum* reproduction is at ∼8 weeks of age, animals of this age are considered as adult *D. cerebrum*. We primarily used male animals for experiments, though some female animals of comparable size were used in group experiments.

Zebrafish and *D. cerebrum* older than 28 days old were tested in a transparent acrylic 300 mm circular arena (black walls and transparent bottom lined with white styrene to diffuse light) filled with 1 L of system water, covered with black curtain to control light and dark. *D. cerebrum* younger than 28 days old fish were placed in a 150 mm petri dish placed in the middle of the aforementioned circular arena filled with 150 mL of system water, in order to allow for sufficient tracking despite their small size. The arena was illuminated with bottom-projected white light (AnyBeam Pico Projector, HD301M1-H2) and infrared light (CM-IR130-850NM, CMVision Technologies Inc.) Behavioral videos were acquired from a camera mounted above the arena, at 121 Hz (FLIR Grasshopper, #33-534) with an 8 mm/ F 1.8 lens (Edmund Optics, #15-626) and long-pass filter (Edmund Optics, #12-767).

Before the recording, fish were left to habituate in the arena for 10 min, and then recorded for 5 min. In light-dark experiments, fish were recorded under light for 10 min, then lights were turned off and fish were recorded for an additional 15 min. Since fish are affected by light-dark transitions (*44*), we considered the first 10 min of dark as habituation and only the last 5 min were analyzed. We note that the 8 week-old fish used in the light-dark experiments were also included in the 8 week-old group in the development experiments. For clear-wall assays, the 300 mm arena was divided by a 1 mm-thick transparent acrylic barrier sealed to the wall and bottom of the arena to prevent water exchange.

#### *In vivo* two-photon calcium imaging of head-tethered *D. cerebrum*

For live *D. cerebrum* brain imaging, we designed a triagonal chamber filled with fish housing water, built from two LCD screens (HAMTYSAN, 7 inch 800×480 LCD) and a transparent acrylic back wall and floor for IR lights and recording camera, respectively. A 4.5 mm column of clear acrylic was used as a pedestal to position head-fixed fish at a standard positioning within this arena: the arena was 10 cm high in total, with the fish placed at the intersection of each screen’s midpoint. The two screens occupied ∼260 ° of visual space in azimuth (130 ° each side, with ∼10 ° gap in the front, and ∼90 ° gap behind the fish), and ∼105 ° of visual space in elevation (52.5 ° below and above, though note that water height only reached ∼3 mm above the fish’s eye).

To image the brains of *D. cerebrum* tethered in this environment, we developed a head-fixation protocol (see **fig. S5A**). We prepared a 24×30×1 mm coverslip (12545B, Fisher Scientific) by lining UV-cured plastic (Bondic) on the margins to hold agarose. We then placed an anesthetized *D. cerebrum* (anesthetized in 120 mg/L MS-222 for 1-2 minutes) upside down by the front edge of the coverslip, secured its position with two wedges of pre-cured Sylgard placed behind each ear, and covered it with 3-4% low-melting point agarose (UltraPure^TM^ LMP Agarose, Invitrogen) – ensuring the back of the head and trunk are as close as possible to the glass. We removed the agarose covering the mouth, gills, and tip of the tail with a scalpel, and ventilated with oxygenated water to ensure gill movement. We then applied a strip of UV-curable plastic extending from each side of the coverslip, including the agarose-covered part of the fish, to provide extra stability. UV light was delivered for less than a second, and the mounted fish was placed in fresh fish water for recovery. Fish that showed good recovery by moving their tail and ventilating their gills normally were transferred into the chamber described above (the success rate of this preparation was approximately 70% for 2 weeks old fish, 50% for 4 weeks old and 30% for 6 weeks old). Coverslips were suspended from the acrylic pedestal using 3 × 1 mm magnets (Ethcool): two glued into the top of the pedestal, and two on top of the coverslip. The arena was then filled with warmed (28 °C) and oxygenated system water (∼ 500 mL), and fish were allowed to recover for 10-60 minutes before imaging data were collected.

For tail tracking, fish were illuminated from behind by two collimated 850 nm LEDs (ThorLabs, M850L3), and tail movements reflected from a mirror beneath the chamber were recorded by a 121 Hz IR camera (FLIR Grasshopper, #33-534) with a 25 mm/ F 1.85 lens (Edmund Optics) and IR bandpass filter (Thorlabs, FBH850-40). BonsaiRx was used to track tail movements online, measure the timing of microscope frame acquisitions, and display visual stimuli. Visual stimuli were displayed on the screens using cube mapping in the BonVision package (*42*), where each screen displayed a viewport in a virtual environment. Global motion stimuli were generated by producing an image of evenly spaced red dots on a black background, and tiling this image along the inside of a sphere. The observer was placed at the center of this sphere in the virtual environment, and the sphere rotates in the y axis either left or right at 20 °/s, reversing the direction of rotation every 20 s.

In the case of social motion stimuli, we extracted a short 5 s period of schooling behavior from 121 Hz recordings of a group of 4 eight-week-old *D. cerebrum*, and obtained the centroid position and heading direction of each fish. We selected one of the fish as the “focal” individual, and translated/rotated all points around this focal individual, to produce a time series of each fish’s position and orientation in reference to the focal individual (which was held at the position of 0,0 and heading angle of 0; see **Fig. 3A**). A 2 s period where no other fish were visible was appended to the end of the stimulus. We subsampled this 7 s dataset of 3 fish (“virtual group members”) to 60 Hz in order to accommodate our screens’ refresh rate, and saved the egocentric centroids and heading angles of each fish as a .csv file. In BonsaiRx, we developed a virtual environment, composed of a 1500 mm^2^ plane with a smoothed white-noise pattern (“seafloor”). The fish’s position in this VR world was in the center (0,0 in x/y), 20 mm above the seafloor. Virtual group members objects were presented 10 mm above the focal fish, with shapes created as ellipsoids with a scale of 8×1×3 mm in x/y/z (horizontal stimulus), 3×1×8 mm in x/y/z (vertical stimulus), or 3×3×3 mm in x/y/z (spherical stimulus), and moved according to the x/y/angle data provided for each of the 3 fish in the field of view. For horizontal stimuli, which were polarized in azimuth, the direction of the shapes moved according to the measured heading direction angles. After a 120 s baseline period where only the seafloor was visible, we used a trial structure to present these 7 s stimuli to head-tethered fish, with 9-11 s inter-trial intervals, and random selection of these three stimulus shapes or the x/y movement patterns of one fish applied to the seafloor (“background” social motion stimulus). All stimulus environments were displayed in shades of red to prevent interference with GCaMP imaging.

Two-photon microscopy was performed with a ThorLabs Bergamo II multiphoton microscope controlled by ThorImageLS 4.1 and illuminated by an fs-pulsed 80-MHz Ti:S laser (MaiTai DeepSee; SpectraPhysics). GCaMP6s fluorescence was imaged with a 16x / 0.8W Nikon objective at an excitation wavelength of 930 nm and ≤10 mW excitation power. Frames of 800 × 800 pixels covering a field-of-view of 833×833 um, were acquired at a rate of 19.3 Hz, and averaged three times to produce a final effective imaging rate of 6.43 Hz. While imaging fish across different ages, we focused on single z-planes that maximized the number of neurons visible in the telencephalon and mesencephalon, while including some neurons in the diencephalon (primarily habenula and dorsal thalamus), and the hindbrain (though note that more of the hindbrain was visible in the smaller two-week-old fish). We imaged fish at a diagonal, to include as much of the brain as possible in older fish, which have a wider midbrain and elongated forebrain (*19*).

#### Behavior Analysis

We used Social LEAP-Estimates Animal Poses (SLEAP; (*27*)), for tracking the location and posture of fish in groups. Each video was checked and proofread after pose inference and identity tracking, and output files (.h5) containing the locations of each point were further analyzed using python code organized in Colab notebooks (available upon publication). The SLEAP model was trained to track each fish with six points distributed from the nose to the proximal tail, while we excluded the most distal tail owing to the limited opacity of this structure. For each fish at each time point, the animal centroid was defined as the x/y coordinate labeling the front of the swim bladder, the nose was defined as the midpoint between the x/y coordinates of each eye, and the tail tip was defined as the furthest reliable x/y coordinate along the tail (second-to-last point). Heading angle was calculated as the direction following the vector between the centroid and nose coordinates. All x/y coordinates were transformed from pixels to mm for subsequent quantification. The time series of x/y positions and heading angles were smoothed by a five-frame running average (equaling 60 ms time windows). Speed (in x/y position, or in angle) was calculated from the change in position or heading angle across frames, and was subsequently smoothed by a ten-frame running average (120 ms). Nearest neighbors were identified for each fish at each time point, and the angle between a focal fish and their nearest neighbor was identified as the absolute difference in heading angle.

Fraction time moving was calculated as the number of frames where fish moved > 2 body length/s. Heading angle correlation is the average pairwise Pearson’s correlation coefficient between all fish pairs, over the full length of the recording. Group area was defined as the convex hull in two dimensions, from all the points across all fish in the group. For experiments analyzing social interactions across a clear barrier, we only examined fish pairs where the mean inter-animal distance is under 25 mm across the recordings, and only included frames where fish are within the middle 80% of the wall (avoiding the 10% closest to each arena boundary). We identified the difference in heading angle as the absolute value of the heading angle difference between closely-spaced individuals.

Egocentric spatial maps were calculated for each individual fish, and averaged across every 10^th^ time point for all fish in the group. For occupancy maps, the focal fish’s centroid was placed in the center of an empty 100 × 100 array of zeros (covering 100 mm^2^ or 6 body length^2^), and oriented to face north in every frame. All other fish centroids were displaced and rotated around the focal fish. If another fish was within the 100 × 100 array, that pixel was assigned a 1. This array was produced for each time point for a focal fish, then down-sampled to 50 × 50 and summed to produce a final 50 × 50 array. This same process was used to calculate speed maps, but instead of pixels being assigned a value of 1 when a fish was present in that location, the value was assigned as the linear speed (x/y displacement from the previous frame) or angular speed (degree change from the previous frame) of the focal fish.

#### Imaging Analysis

Before commencing analysis, we only proceeded with experiments where fish showed spontaneous tail movement but did not show substantial z-movements. Imaging data were motion corrected in Suite2p (*45*), followed by cell extraction. Time series were inspected to ensure there was no z-motion contamination or slow drift, and only included neurons with continuous measurements and which were classified as a cell by Suite2p (“iscell”=1). We excluded the first 5% of each imaging trial to avoid the influence of sound-evoked activity upon the initiation of scanning. Fluorescence traces were detrended, smoothed with a 1 s rolling mean, and z-scored. Neurons in each recorded fish were further classified as belonging to the telencephalon, diencephalon, mesencephalon, and rhombencephalon with manually-defined boundaries. Z-scored fluorescence and the median x/y coordinates for each cell within these boundaries were used for further analysis. Final counts of neurons included (N=4 fish in each age) are: telencephalon (forebrain; N = 734 cells at two weeks, 1016 cells at four weeks, 1669 cells at six weeks), diencephalon (N = 130 cells at two weeks, 199 cells at four weeks, 435 cells at six weeks), mesencephalon (midbrain; N = 1580 cells at two weeks, 1672 cells at four weeks, 1726 cells at six weeks), and rhombencephalon (hindbrain; N = 980 cells at two weeks, 381 cells at four weeks, 242 cells at six weeks). We quantified responsive cells as neurons whose maximum z-scored fluorescence response to any social motion object stimulus (averaged across trials) was greater than 2.5 standard deviations above the mean.

All analyses were conducted using python code organized in Colab notebooks (available upon publication). For behavioral analysis of tethered fish, we smoothed mean-subtracted tail angles, and analyzed the sum of the tail angle’s absolute value over the 7 s stimulus for each trial. This value was divided by the mean tail angle sum across five pre-stimulus periods of the same length. Values below 1 indicate less stimulus-driven movement than baseline movements, whereas values above 1 indicate more. For neural activity analysis, behavioral recordings (tail movement, stimulus presentation) were first cropped to align with the times of two-photon image acquisition, and time series were resampled to a common 10 Hz sampling rate for further analysis. To exclude low-amplitude tail movements, we only analyzed tail movements that exceeded 15° from the midline, and produced an array of binary movement times for subsequent analysis.

We fit linear models to each neuron using ridge regression (alphas = 0.001, 0.01, 0.1, 1.0, 10.0), training on the first 75% of the data and testing on the last 25%. Regressors were normalized between 0 and 1, and smoothed by an exponentially weighted moving average with decay of 3 s (approximate decay of H2B-GCaMP6s). For optic flow stimuli, the regressors were the direction of the global optic flow (accumulating positive to go right, and negative to go left) and tail movements. For social motion stimuli, regressors were boxcar functions for each stimulus type (horizontal shape, vertical shape, spherical shape, and background motion), and tail movements. We used the model coefficients (βs) for each regressor to identify neurons in each age group with the most selective responses to each stimulus, and plotted these neurons’ averaged peri-stimulus activity. We defined the “most responsive cells” of each age group to be those in the top 5^th^ percentile of the β distribution for a given regressor type. We quantified a neuron’s stimulus selectivity by measuring the absolute value of the difference between the maximum z-scored activity (σ) during the 7 s of the horizontal stimulus (averaged across all trials), and the maximum z-scored activity (σ) during the 7 s of the vertical or background stimulus (averaged across all trials).

To analyze population-level activity, we performed principal components analysis (PCA) on populations of neurons in each fish with a positive prediction score from the linear model. These traces were mean-centered before PCA; 15 components explained ∼ 50-80% of the variance for each fish. For analysis of subsampled populations, we randomly selected 40 cells from the original distribution, before PCA (with the exception of one two-week-old fish with 18 cells and one four-week old fish with 32 cells). For visualization, we plotted the trial-averaged trajectories in the first 3 components. To quantify the distance between trajectories in 15-dimensional PC space, we obtained the mean pairwise distances amongst all 3 object trajectories or the mean pairwise distances between each object trajectory and the moving floor stimulus, over the 7 s stimulus presentation period. We used both cosine distance (one minus cosine similarity) and Euclidean distance as metrics. Displayed time series of cosine distance or Euclidean distance are baseline-subtracted for display.

#### Statistics

Groups were tested for normality using the Shapiro-Wilk test for normality. Non-parametric tests (Mann-Whitney U, Kruskal-Wallis H) were conducted if Shapiro-Wilk *P*<0.05 for any group or if groups included measurements of each individual from a group. Otherwise, parametric tests were used. *Post hoc* tests were conducted if the main effect was *P*<0.05, except in the case of comparing model coefficients across regressors, where we used a Bonferroni correction to correct for multiple Mann-Whitney U tests across 5 or 2 groups, and therefore set significance cutoff to *P*<0.01, or *P*<0.025, respectively. *Post hoc* tests (Dunn’s test for non-parametric data, or *t*-tests for parametric data) were all corrected for multiple comparisons with a Bonferroni correction. Exact tests and *P* values are reported in the figure legends.

**Fig. S1.**
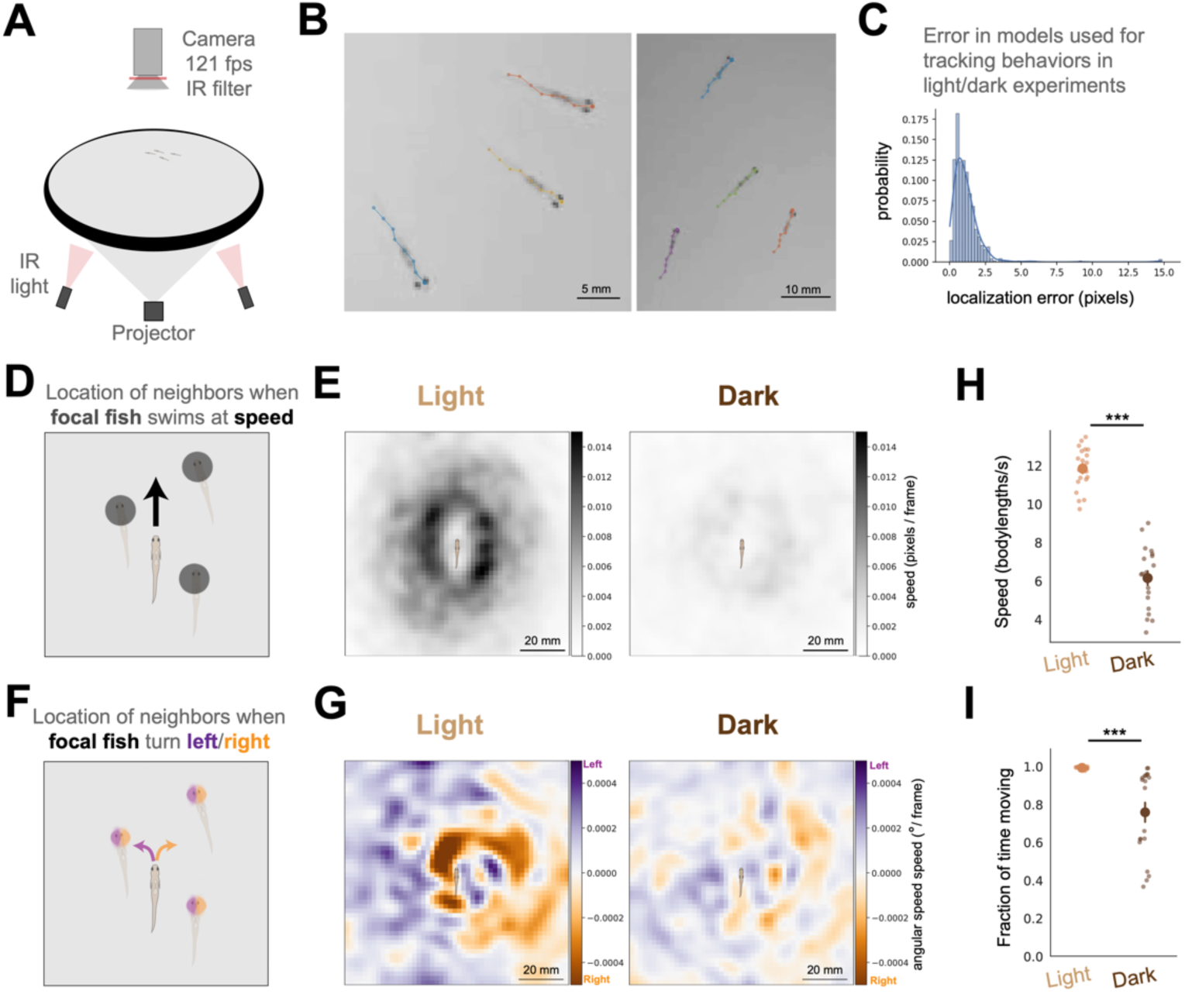
Additional descriptions of *D. cerebrum* schooling behavior. A) Schematic of recording configuration. B) Example frames of animals labeled by SLEAP. C) Measurement of tracking error (pixels) in model used to track fish in light and dark experiments. D) Schematic of speed measurement: speed of focal fish, relative to the position of other individuals. Schematic of focal fish is shown (approximate body size). E) Egocentric speed heatmaps, for groups of four fish swimming in light (left), or dark (right). Darker colors indicate higher speed of focal fish (pixels/frame) when other fish occupy the positions in the heatmap. N = 5 groups, 121 frames/s recordings for 300 s each in light and dark. Average of all timepoints across all fish. Schematic of focal fish is shown (approximate body size). F) Schematic of angular speed measurement: angular speed of focal fish, relative to the position of other individuals. G) Egocentric angular speed heatmaps, for groups of four fish swimming in light (left), or dark (right). Purple colors indicate faster leftward rotation speed (°/frame) and orange indicates faster rightward rotation speed of focal fish when other fish occupy the positions in the heatmap. N = 5 groups, 121 frames/s recordings for 300 s each in light and dark. Average of all timepoints across all fish. Schematic of focal fish is shown (approximate body size). H) Mean speed (body lengths/s). comparing between groups of four fish swimming in light versus dark (N = 20 fish from 5 groups). Mann-Whitney U = 400.0. *P* = 6.7956 × 10^-8^. I) Mean fraction of time spent moving (>2 body lengths/s) comparing between groups of four fish swimming in light versus dark (N = 20 fish from 5 groups). Mann-Whitney U = 386.0. *P* = 5.2269 × 10^-7^. \*\*\**P* < 0.0001.

**Fig. S2.**
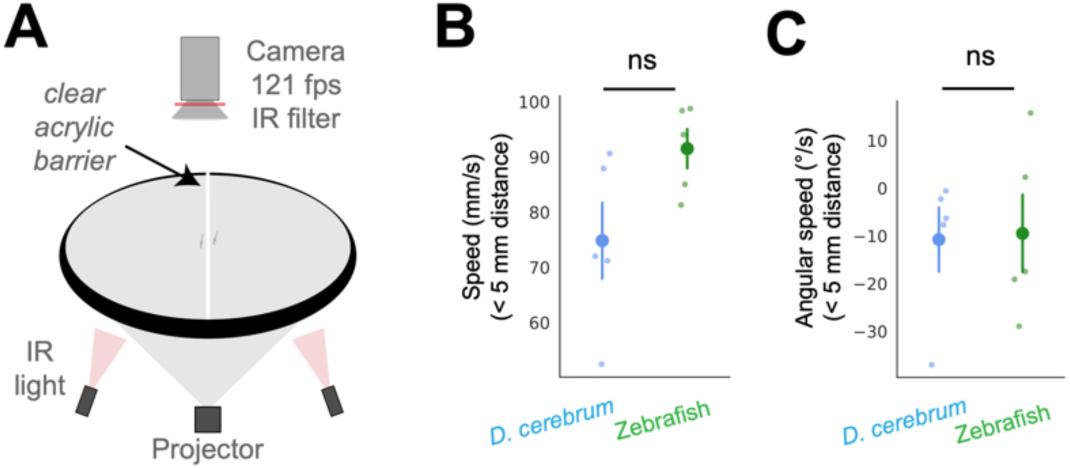
Additional measures of interactions across a clear wall. A) Schematic of recording configuration, with clear water-tight barrier between fish. B) Mean speed (mm/s) for fish within 5 mm of each other across the barrier. Mann-Whitney U = 4.0. *P* = 0.0952. C) Mean angular speed (°/s) for fish within 5 mm of each other across the barrier. Mann-Whitney U = 12.0 *P* = 1.0.

**Fig. S3.**
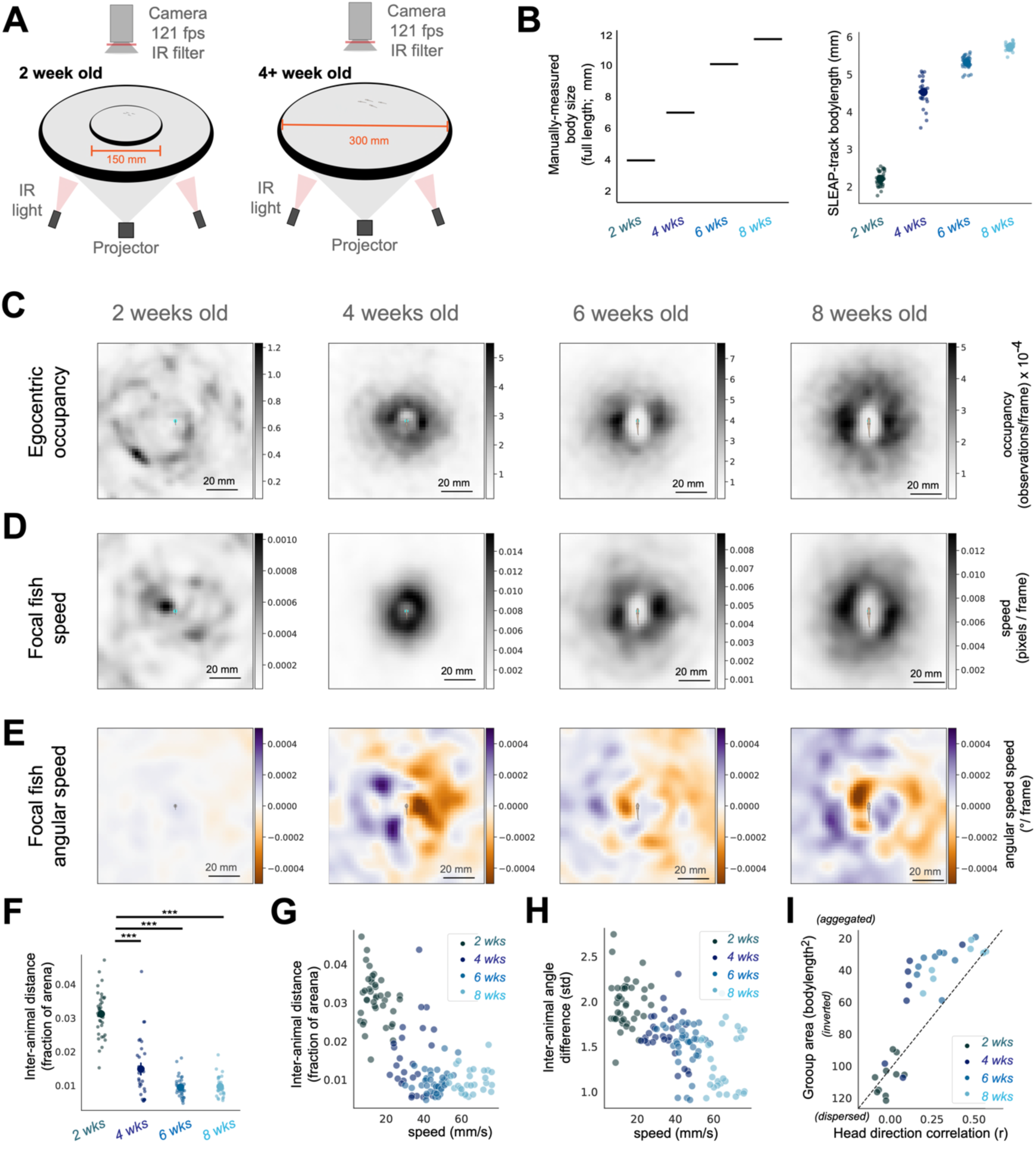
Additional descriptions of *D. cerebrum* behavioral maturation. A) Schematic of recording configurations: 150 mm diameter arena for 2-week old fish, and 300 mm diameter for fish 4-weeks old and older. B) Measurements of 2-, 4-, 6-, and 8-week old *D. cerebrum* body length: manual (left) and extracted from SLEAP tracking (right). Note that SLEAP tracking reliably tracks the most visible body trunk of the *D. cerebrum.*, while omitting the nearly-transparent distal tail (and thus tracks a shorter length). C) Egocentric occupancy heatmaps, for groups of four fish (left to right: 2, 4, 6, or 8 weeks old). Maps are not scaled by body length, in contrast to body length-scaled data in Figure 2C. Average of all timepoints across all fish. Schematic of focal fish is shown (approximate body size). D) Egocentric speed heatmaps, for groups of four fish (left to right: 2, 4, 6, or 8 weeks old). Maps are not scaled by body length. Average of all timepoints across all fish. Schematic of focal fish is shown (approximate body size). E) Egocentric angular speed heatmaps, for groups of four fish (left to right: 2, 4, 6, or 8 weeks old). Maps are not scaled by body length. Average of all timepoints across all fish. Schematic of focal fish is shown (approximate body size). F) Mean inter-animal distance (fraction of arena size) comparing groups of four fish. Kruskal-Wallis followed by *post hoc* Dunn’s test with Bonferroni correction. H = 79.666, *P* = 3.6189 × 10^-17^. G) Scatterplot of mean speed (mm/s) and mean inter-animal distance (fraction of arena), with each fish colored by age. H) Scatterplot of mean speed (mm/s) and mean standard deviation of the inter-animal angle difference, with each fish colored by age. I) Scatterplot of mean group area (bodylengths^2^) and mean head direction correlation (r), with each group colored by age. Dotted line indicates expectation from linear relationship. N = 10, 7, 9, and 7 groups for 2, 4, 6, and 8 week-old fish, respectively. \*\*\**P* < 0.0001.

**Fig. S4.**
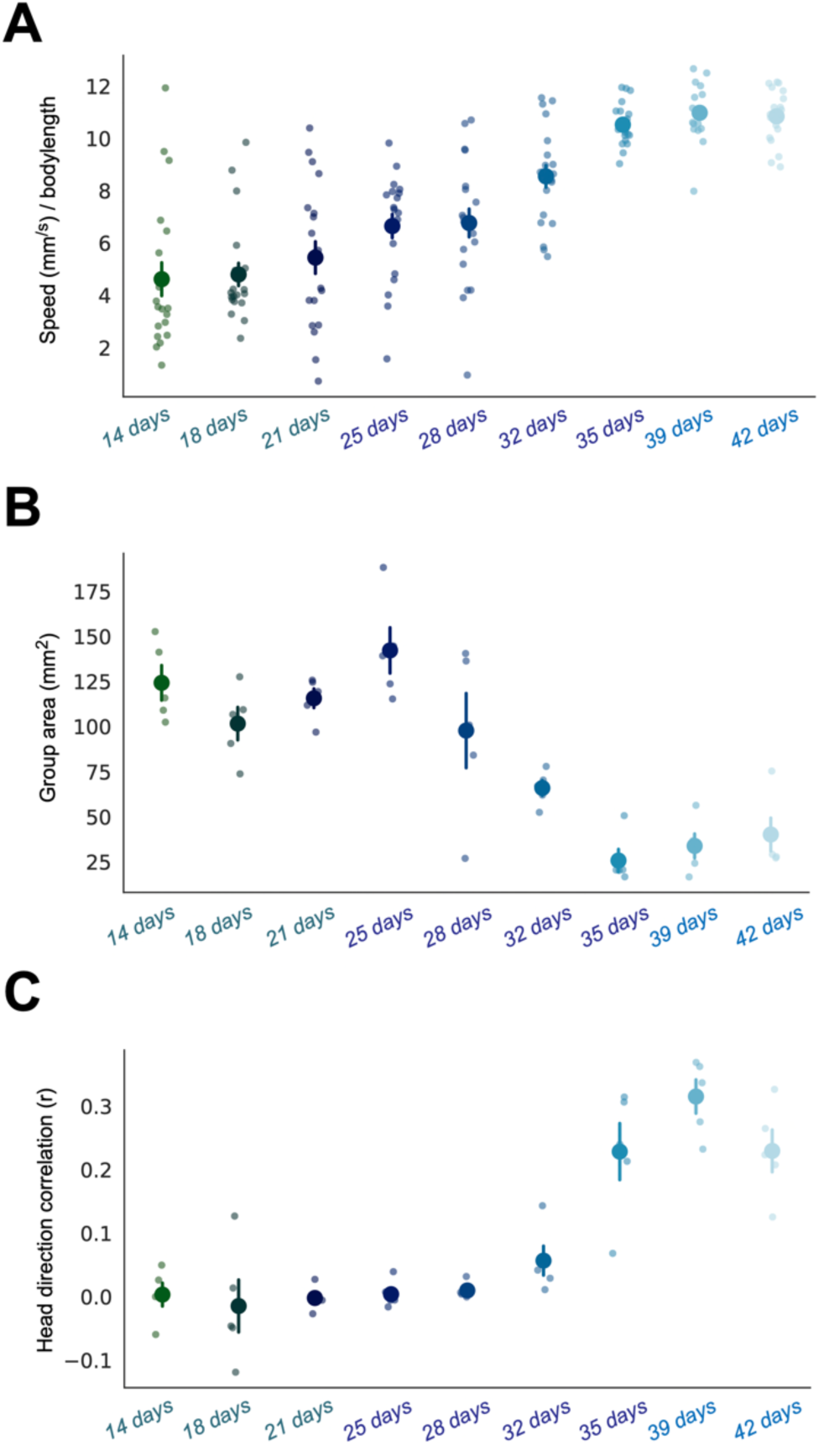
Longitudinal measurement of social development in *D. cerebrum*. A comparison of groups over 14, 18, 21, 25, 28, 32, 35, 39, and 42 days old – repeatedly testing 5 groups of 4 from the same tank (though the exact same identity of each group was not maintained). A) Mean speed (body lengths/s), comparing each fish in groups of four (N=20 total). Kruskal-Wallis followed by *post hoc* Dunn’s test with Bonferroni correction. H = 134.682, *P* = 3.022 × 10^-25^. B) Mean group area (mm^2^), comparing groups of four fish. Kruskal-Wallis followed by *post hoc* Dunn’s test with Bonferroni correction. H = 33.846, *P* = 4.332 × 10^-5^. C) Mean of pairwise heading direction correlation coefficients (r) amongst all fish, comparing between groups of four fish. Kruskal-Wallis followed by *post hoc* Dunn’s test with Bonferroni correction. H = 33.02, *P* = 6.107 × 10^-5^. N = 5 groups of 4 fish, repeatedly measured at each timepoint. Mean +/- s.e.m. * *P* < 0.05 \*\**P* < 0.01 \*\*\**P* < 10^-4^.

**Fig. S5.**
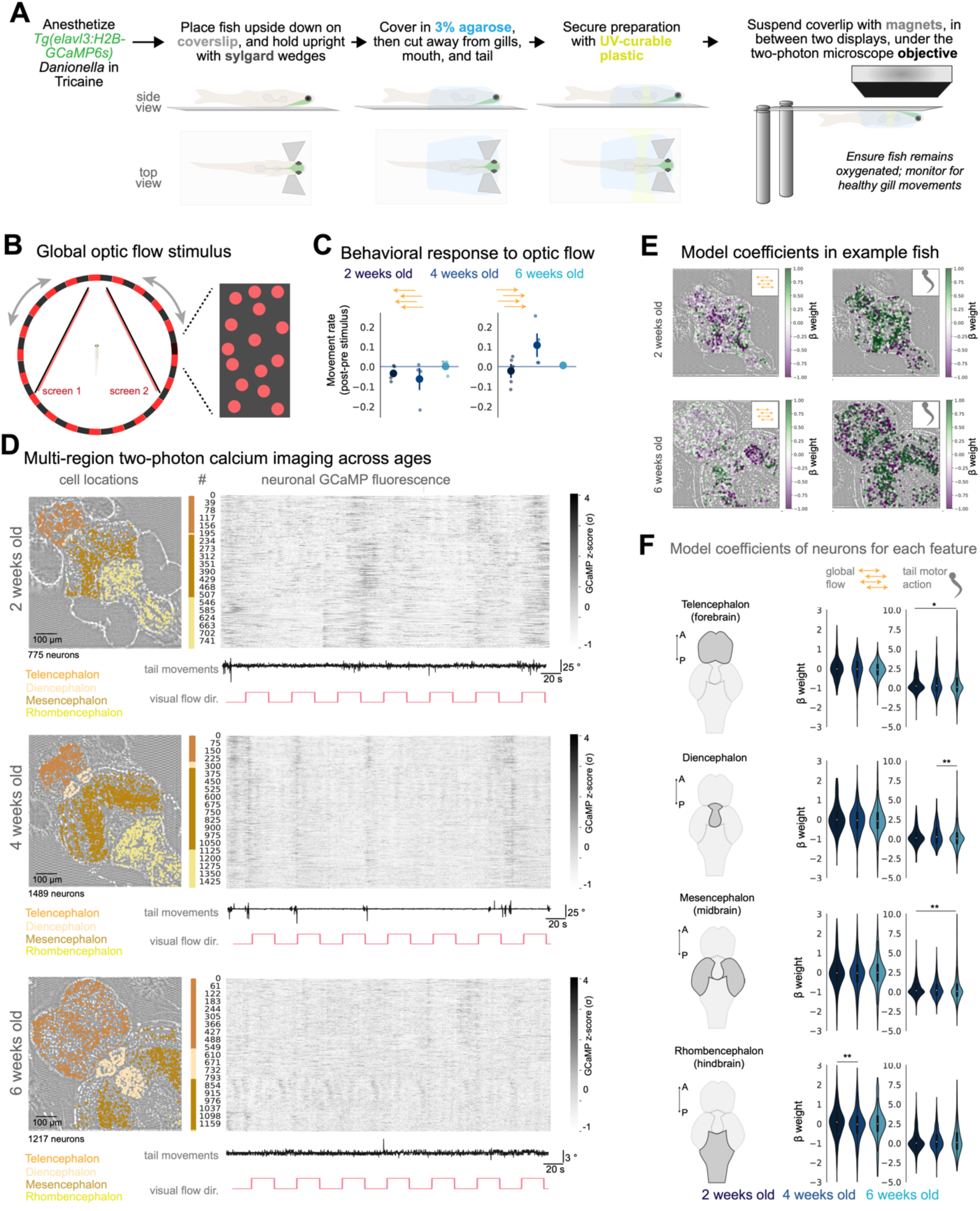
Visuomotor behavior and neural activity in head-tethered *D. cerebrum*. A) Schematic of head-fixation preparation for imaging behaving *Tg(elavl3:H2B-GCaMP6s) D. cerebrum.* All steps are typically completed in under ten minutes, with supplemental oxygenated water provided for the fish between steps. B) Schematic of global optic flow stimulus: Black background with red dots, rendered as a sphere rotating around the fish left and right. C) Behavioral response to global optic flow stimuli. Data is the mean of the post-stimulus movement rate (5 s) - the pre-stimulus movement rate (2 s). D) Example fish of each age, with anatomically-annotated cell locations (left), and z-scored neural activity for all recorded neurons (right). Global optic flow stimulus timing and tail movements are noted below each neural activity heatmap. A subset of the full imaging time is displayed. E) Location of neurons in example fish at 2- and 6-weeks of age, with neurons colored by model coefficients (β) for the global motion stimulus direction (left) or tail movements (right). F) Summary data for all neurons across all ages, separated by gross anatomical subdivision (left; A=Anterior; P=Posterior). We compare model coefficients (β) for each neuron within anatomical boundaries, across the 2 regressors. Kruskal-Wallis H-tests for each regressor are adjusted by a Bonferroni correction for multiple comparisons, across age groups within a regressor type. Only the results of significant adjusted H-tests (*P*<0.025) are reported. N = 5 (Wk2), 4 (Wk4), and 6 (Wk6) separate fish. Telencephalon: N = 638 (Wk2), 1255 (Wk4), 3406 (Wk6) cells. Motion stimulus: H = 3.349, *P* = 0.187. Tail movement: H = 9.588, *P* = 0.0083. Diencephalon: N = 205 (Wk2), 245 (Wk4), 1067 (Wk6) cells. Motion stimulus: H = 3.235, *P* = 0.198. Tail movement: H = 12.343, *P* = 0.0021. Mesencephalon: N = 1874 (Wk2), 2078 (Wk4), 3212 (Wk6) cells. Motion stimulus: H = 5.204, *P* = 0.074. Tail movement: H = 15.37, *P* = 0.00046. Rhombencephalon: N = 1070 (Wk2), 876 (Wk4), 320 (Wk6) cells. Motion stimulus: Kruskal-Wallis H = 14.529, *P* = 0.0007. Tail movement: H = 4.734, *P* = 0.0938. * *P* < 0.05 \*\**P* < 0.01.

**Fig. S6.**
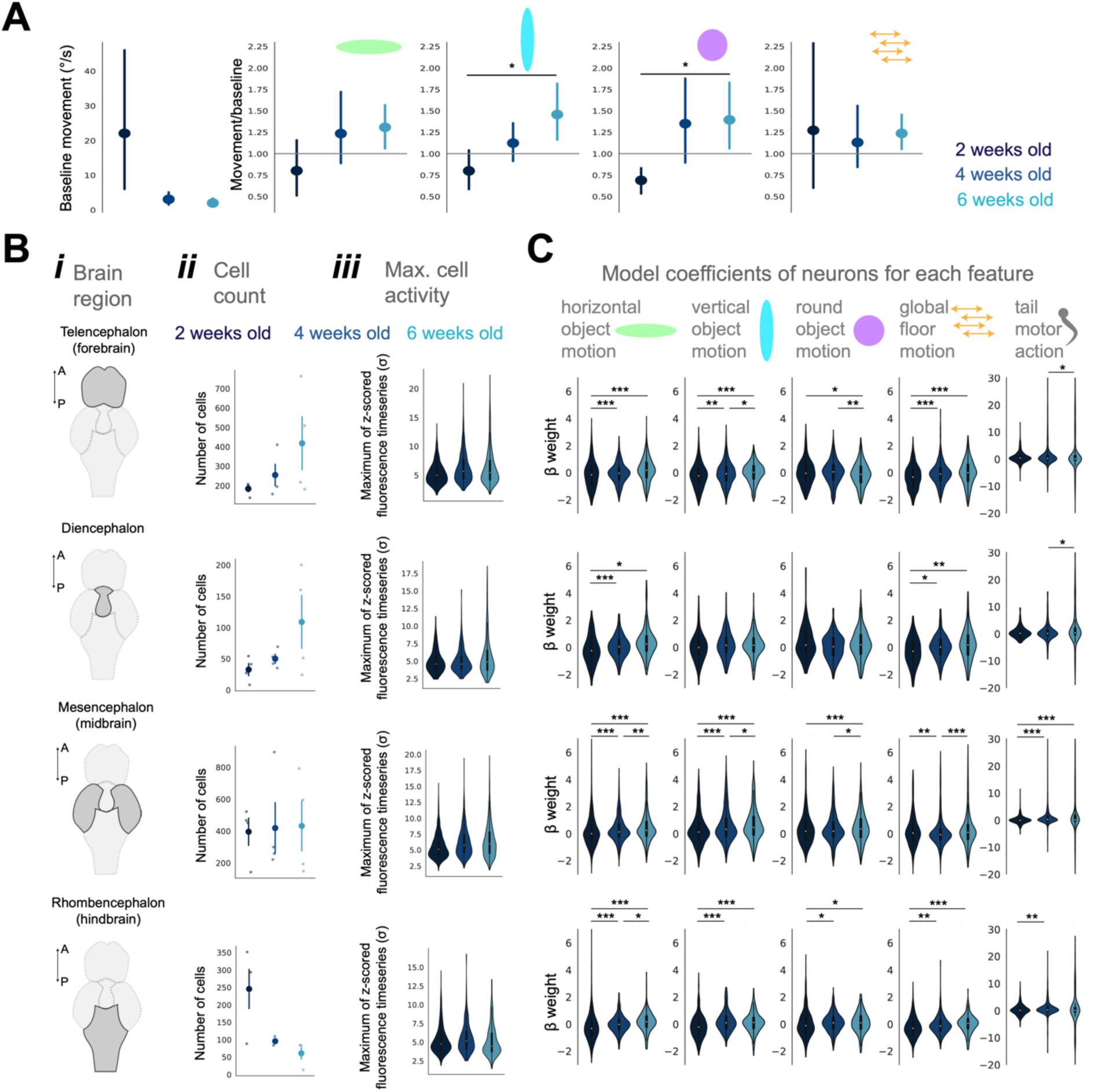
Additional descriptions of neural responses to social motion stimuli. A) Left: baseline movement rate. Right: Behavioral response to social motion stimuli, normalized by baseline rate. Mean +/- 95% confidence intervals are shown. One-way ANOVA followed by *post hoc* t-tests with Bonferroni correction (Horizontal shape: F= 2.2, *P* = 0.11; Vertical shape: F= 5.37, *P* = 0.007. Spherical shape: F= 4.21, *P* = 0.019; Background motion: F = 0.05, *P* = 0.94). B) For each brain region (*i*), displayed is the number of cells per brain region in each age (*ii*), and the maximum z-scores for cells in each age (*iii*). C) Across the brain regions defined in B, summary data from linear models fit to all neurons across all ages are displayed, and separated by anatomical subdivision (left; A=Anterior; P=Posterior). We compare model coefficients (β) for each neuron within anatomical boundaries, across the 5 regressors. Kruskal-Wallis H-tests for each regressor are adjusted by a Bonferroni correction for multiple comparisons, as is each pairwise *post hoc* Dunn’s test, across age groups within a regressor type. Only the results of significant adjusted H-tests (*P*<0.01) are reported. N = 4 (Wk2), 4 (Wk4), and 4 (Wk6) separate fish. From top: Telencephalon: N = 734 (Wk2), 1016 (Wk4), 1669 (Wk6) cells. Horizontal shape: H = 108.99, *P* = 2.157 × 10^-24^. Vertical shape: H = 50.242, *P* = 1.231 × 10^-1^ . Spherical shape: H = 21.925, *P* = 1.734 × 10^-5^. Background motion: H = 64.571, *P* = 9.509 × 10^-15^. Tail movement: Kruskal-Wallis H = 11.749, *P* = 0.00281. Diencephalon: N = 130 (Wk2), 199 (Wk4), 435 (Wk6) cells. Horizontal shape: H = 36.685, *P* = 1.081 × 10^-8^. Vertical shape: H = 6.388 *P* = 0.041. Spherical shape: H = 5.553 *P* = 0.062. Background motion: H = 19.389, *P* = 6.161 × 10^-5^. Tail movement: H = 10.392, *P* = 0.0055. Mesencephalon: N = 1580 (Wk2), 1672 (Wk4), 1726 (Wk6) cells. Horizontal shape: H = 94.146, *P* = 3.601 × 10^-21^. Vertical shape: H = 107.914, *P* = 3.689 × 10^-24^. Spherical shape: H = 19.799, *P* = 5.018 × 10^-5^. Background motion: H = 29.210, *P* = 4.541 × 10^-7^. Tail movement: H = 162.349, *P* = 5.577 × 10^-36^. Rhombencephalon: N = 980 (Wk2), 381 (Wk4), 242 (Wk6) cells. Horizontal shape: H = 91.549, *P* = 1.319 × 10^-20^. Vertical shape: H = 84.639, *P* = 4.177 × 10^-19^. Spherical shape: H = 17.757, *P* = 1.394 × 10^-4^. Background motion: H = 47.971, *P* = 3.829 × 10^-11^. Tail movement: H = 14.995, *P* = 5.544 × 10^-4^. * *P* < 0.05 \*\**P* < 0.01 \*\*\**P* < 10^-6^

**Fig. S7.**
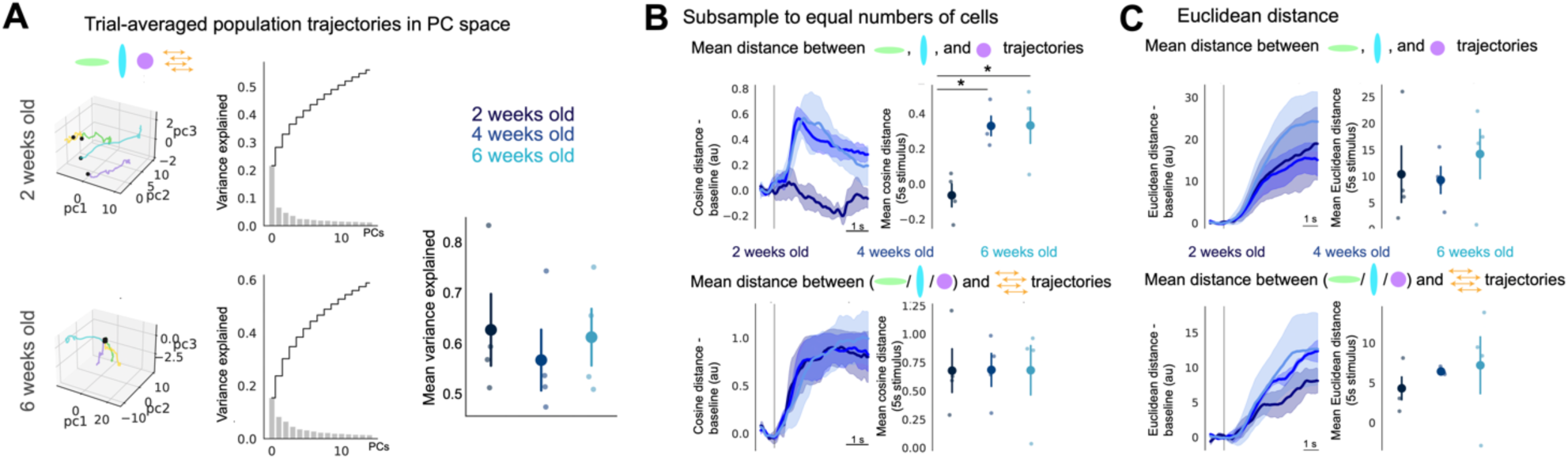
Low-dimensional neural responses to social motion stimuli. A) Left: low-dimensional trajectory of trial-averaged social motion stimulus responses in two example fish (different example fish than in Fig. 4), beginning at stimulus onset (black dot) and progressing for 7 s post-stimulus. Trajectories are colored by stimulus type, and are shown for the first three principal components (pcs). Right: mean variance explained by 15 PCs used for analysis (One-way ANOVA F = 0.24, *P* = 0.78). B) Top: Mean pairwise cosine distance of trial-averaged trajectories in 15-dimensional PC space, between each of the three virtual group member social motion trial types (horizontal, vertical, or spherical shape). Neurons from each fish have been subsampled to be equal in number across each fish (see *Methods*). Left: Timeseries are baseline-subtracted (1 s before stimulus onset), and mean +/- s.e.m is shown. Right: mean across the 7 s stimulus period. One-way ANOVA F = 8.88, *P* = 0.007. *Post hoc* t-test: Wk2 vs Wk4: *P* = 0.003. Wk2 vs Wk6: *P* = 0.016. Bottom: Mean pairwise cosine distance of trial-averaged trajectories in 15-dimensional PC space, between each of the three virtual group member social motion trial types (horizontal, vertical, or spherical shape) and the background motion trials. Neurons from each fish have been subsampled to be equal in number across each fish. Left: Timeseries are baseline-subtracted (1 s before stimulus onset), and mean +/- s.e.m is shown. Right: mean across the 7 s stimulus period. One-way ANOVA F = 0.003, *P* = 0.99. C) Top: Mean pairwise Euclidean distance of trial-averaged trajectories in 15-dimensional PC space, between each of the three virtual group member social motion trial types (horizontal, vertical, or spherical shape). Left: Timeseries are baseline-subtracted (1 s before stimulus onset), and mean +/- s.e.m is shown. Right: mean across the 7 s stimulus period. One-way ANOVA F = 0.456, *P* = 0.647. Bottom: Mean pairwise Euclidean distance of trial-averaged trajectories in 15-dimensional PC space, between each of the three virtual group member social motion trial types (horizontal, vertical, or spherical shape) and the background motion trials. Left: Timeseries are baseline-subtracted (1 s before stimulus onset), and mean +/- s.e.m is shown. Right: mean across the 7 s stimulus period. One-way ANOVA F = 0.355, *P* = 0.71. N=4 fish for all groups. * *P* < 0.05

**Fig. S8.**
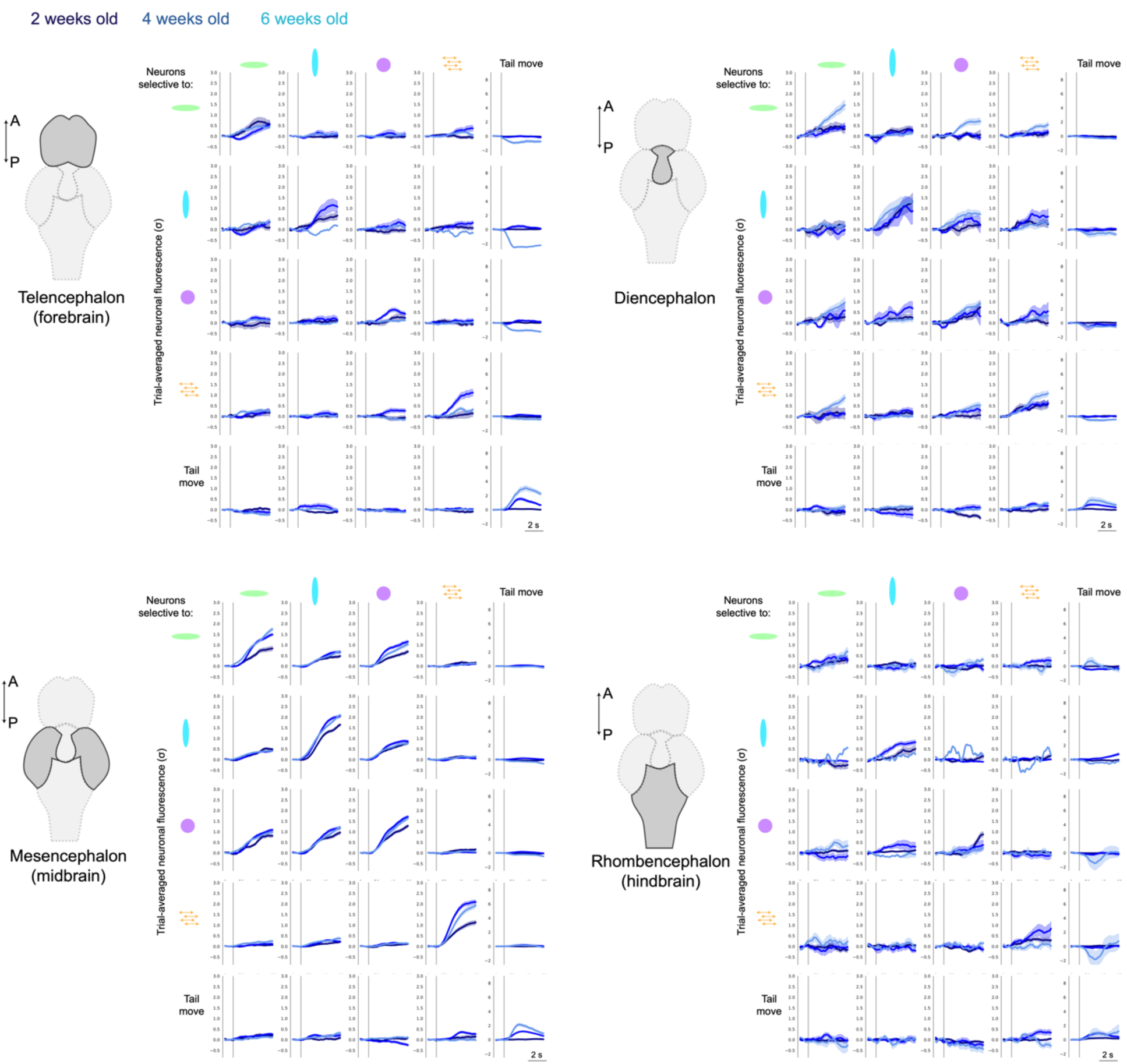
Trial-averaged responses of neurons sorted by model coefficients. Neurons for each age and brain region were selected if their model coefficients was the top 5^th^ percentile of the β distribution (ordered in rows). Their trial-averaged responses are plotted for each stimuli (columns). Traces are mean +/- s.e.m.

**Fig. S9.**
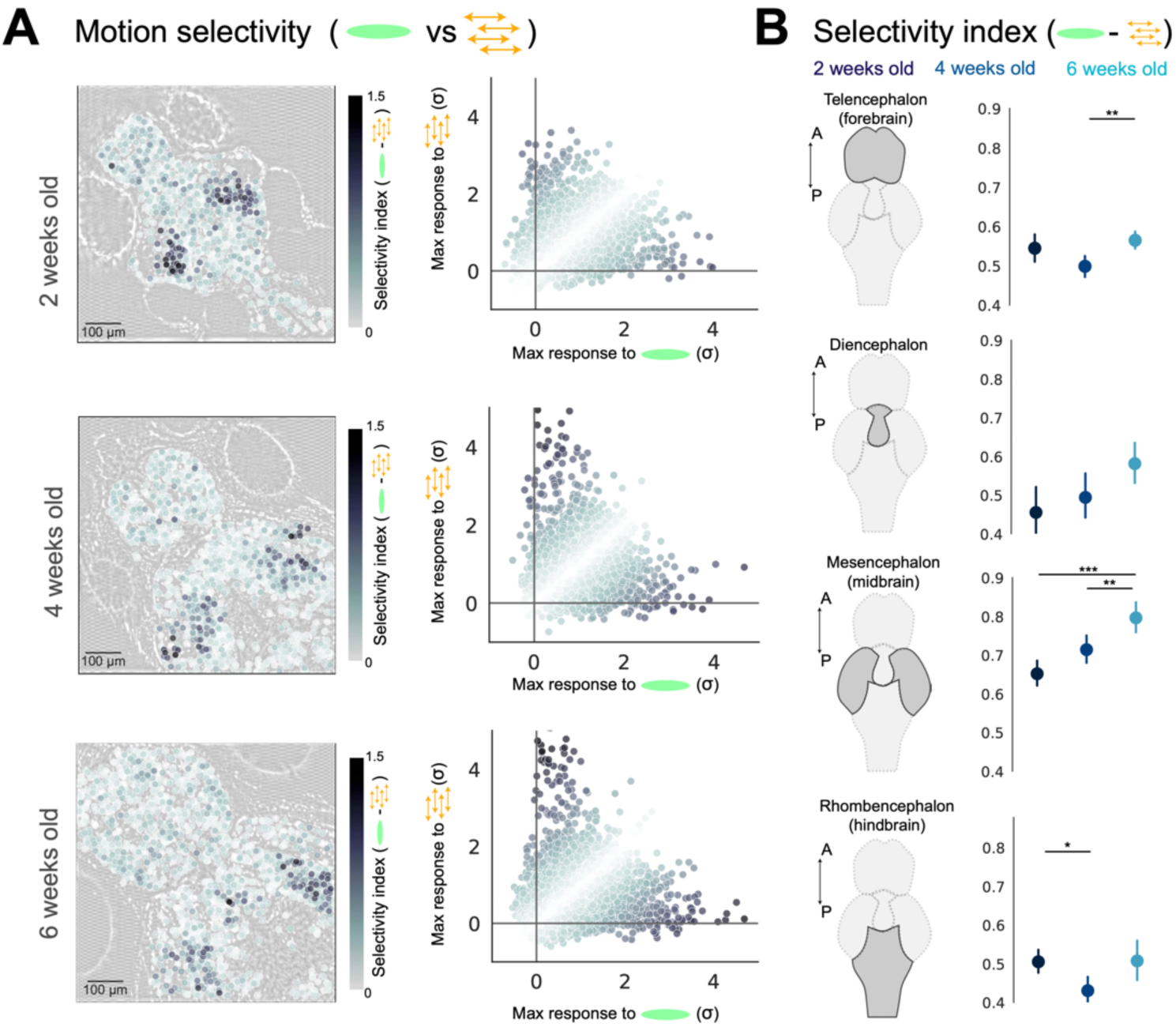
Selectivity of neurons to social motion shape versus background. A) Motion selectivity (absolute value of the difference in the maximum response to horizontal shape and background motion) across each age, displayed on cell locations in example fish (left) and on a scatter plot of stimulus responses across all fish (right; N= 3424 (Wk2), 3268 (Wk4), and 4072 (Wk6) cells.). B) Summary of shape selectivity measure (absolute value of the difference in the maximum response to horizontal shape and background motion) across each age, and separated by anatomical subdivision (left; A=Anterior; P=Posterior). Mean +/- 95% confidence intervals are shown. Statistical significance determined by Kruskal-Wallis H-tests followed by *post hoc* Dunn’s test with Bonferroni correction. From top: Telencephalon: N = 734 (Wk2), 1016 (Wk4), 1669 (Wk6) cells. H = 16.881, *P* = 0.00022. Diencephalon: N = 130 (Wk2), 199 (Wk4), 435 (Wk6) cells. H = 4.070, *P* = 0.131. Mesencephalon: N = 1580 (Wk2), 1672 (Wk4), 1726 (Wk6) cells. H = 25.649, *P* = 2.693 × 10^-6^. Rhombencephalon: N = 980 (Wk2), 381 (Wk4), 242 (Wk6) cells. H = 6.421, *P* = 0.040. * *P* < 0.05 \*\**P* < 0.01 \*\*\**P* < 10^-5^

